# Fusing tree-ring and forest inventory data to infer influences on tree growth

**DOI:** 10.1101/097535

**Authors:** Margaret E. K. Evans, Donald A. Falk, Alexis Arizpe, Tyson L. Swetnam, Flurin Babst, Kent E. Holsinger

## Abstract

Better understanding and prediction of tree growth is important because of the many ecosystem services provided by forests and the uncertainty surrounding how forests will respond to anthropogenic climate change. With the ultimate goal of improving models of forest dynamics, here we construct a statistical model that combines complementary data sources – tree-ring and forest inventory data. A Bayesian hierarchical model is used to gain inference on the effects of many factors on tree growth – individual tree size, climate, biophysical conditions, stand-level competitive environment, tree-level canopy status, and forest management treatments – using both diameter at breast height (DBH) and tree-ring data. The model consists of two multiple regression models, one each for the two data sources, linked via a constant of proportionality between coefficients that are found in parallel in the two regressions. The model was applied to a dataset developed at a single, well-studied site in the Jemez Mountains of north-central New Mexico, U. S. A. Inferences from the model included positive effects of seasonal precipitation, wetness index, and height ratio, and negative effects of seasonal temperature, southerly aspect and radiation, and plot basal area. Climatic effects inferred by the model compared well to results from a dendroclimatic analysis. Combining the two data sources did not lead to higher predictive accuracy (using the leave-one-out information criterion, LOOIC), either when there was a large number of increment cores (129) or under a reduced data scenario of 15 increment cores. However, there was a clear advantage, in terms of parameter estimates, to the use of both data sources under the reduced data scenario: DBH remeasurement data for ~500 trees substantially reduced uncertainty about non-climate fixed effects on radial increments. We discuss the kinds of research questions that might be addressed when the high-resolution information on climate effects contained in tree rings are combined with the rich metadata on tree- and stand-level conditions found in forest inventories, including carbon accounting and projection of tree growth and forest dynamics under future climate scenarios.

## Introduction

Improved understanding and prediction of tree growth is important because of the many ecosystem services provided by forests – global climate regulation via carbon sequestration, provisioning of drinking water, flood regulation, erosion control – in addition to the habitat they provide for many species (*i.e*., supporting biodiversity). Climate change is expected to change the geographic distribution of the forest biome, with uncertain consequences with respect to these ecosystem services (Bonan et al. 2008, Fettig et al. 2013). For example, a matter of great scientific uncertainty is whether forests will continue to be a carbon sink (sequestering ~25% of anthropogenic carbon emissions; Friedlingstein et al. 2010, Pan et al. 2011) or become a carbon source in the future (Kurz et al. 2008, Luo et al. 2015). Forests in interior western North America may be at the leading edge of this change – with documented large-scale mortality events driven by warm drought, insect outbreaks, and increasingly large and severe fires (Allen et al. 2010, Falk 2013, Fettig et al. 2013, Williams et al. 2013, Allen et al. 2015, Millar and Stephenson 2015) – and they are predicted to be especially vulnerable to the climate change projected over the course of the 21^st^ century. One model (Jiang et al. 2013) projects a loss of 50% of the needle-leaf evergreen forest in the western United States. Williams et al. (2013) projected that by the 2050’s, average forest drought stress in the southwestern U. S. will be more severe than the most severe drought conditions of the last 1,000 years. Charney et al. (2016) projected reduced growth of up to 70% for interior western North American trees in the second half of the 21^st^ century compared to the first half of the 20^th^ century.

These projections (and others; Rehfeldt et al. 2006 Coops and Waring 2011, Notaro et al. 2012) derive from a diversity data sources and/or modeling approaches, including dynamic vegetation models, tree-ring data, species distribution models, and physiological models, each of which has some inherent limitations (McMahon et al. 2011, Bellard et al. 2012, Bowman et al. 2013, Friend et al. 2014). More reliable prediction and greater insight into the complex of factors that influence individual tree growth could be gained by using multiple sources of data together, with complementary information or sampling design (Evans et al. 2016). Here, we focus on combining tree-ring and forest inventory data, two powerful and complementary data sources. Tree-ring data have annual resolution and decadal- to centennial-length records, which provides considerable information on the influence of climate on tree growth. However, sampling has traditionally been biased towards the most climate-sensitive individuals (for the purpose of reconstruction of past climates), metadata on tree size and stand characteristics are usually missing, and spatial replication is relatively limited (Babst et al. 2014a). National forest inventory programs, such as the U. S. Forest Service’s Forest Inventory and Analysis (FIA; Gillespie et al. 1999), are spatially comprehensive and unbiased with respect to tree size, providing rich forest structure information at the tree and stand level, but in the western U. S., censuses of individual tree growth (“remeasurement”) are conducted on average once every 10 years.

Precedents for using tree-ring and forest inventory data in parallel or in combination were set by Biondi (1999) and Clark et al. (2007). Biondi (1999) analyzed tree-ring and diameter at breast height (DBH) remeasurement data in parallel at the Gus Pearson Research Natural Area in northern Arizona, to better understand how fire suppression and subsequent high-density recruitment affected the growth of large trees. Clark et al. (2007) laid the groundwork for formal statistical fusion of tree-ring and forest inventory data using a hidden process model, in which the two data types were treated as observations (with error) of the true, underlying (hidden) process of individual tree growth. Thus the model consisted of two measurement error models and a process model. However, their process model did not include fixed effects: tree growth was modeled as a function of year random effects and individual random effects. Individual tree growth is known to be influenced by many factors, including first and foremost tree size, followed by a suite of other factors varying at either the tree-level or the stand- (or site-) level, such as stand basal area, various aspects of climate in multiple months or seasons, disturbances, substrate, etc. (Cook and Briffa 1990, Paine et al. 2011, Bowman et al. 2012). In other words, the problem of understanding individual tree growth is a complex one.

Here we develop a hierarchical Bayesian model that can infer the influence of many factors on individual tree growth, relying on a combination of tree-ring and forest inventory-type (diameter remeasurement) data at once. Hierarchical Bayesian modeling is appropriate for the task because it allows one to formally combine information from multiple data sources, while accommodating complexity and scale (Clark 2005). In this application specifically, it allows us to tailor a statistical model to the structure of the data, including the different levels (tree- vs. stand-) at which predictors are available, and the nature of the relationship between the two types of observations. Further, hierarchical Bayesian modeling allows us to examine variation in radial and diameter increments in terms of conditional probabilities – e.g., the influence of climate conditional upon tree size. We develop this model at a single exceptionally well-studied site, where much is known about the history of disturbance, forest management, climate limitations, phenology, substrate, *etc*. This well-studied site provides the opportunity to unravel a complex multiple regression model problem with confidence. We test hypotheses about the factors influencing individual tree growth using a multiple regression approach, with several main effects, expected nonlinearities, and potential interactions among factors, to evaluate the hypotheses that growth is influenced by tree size, climate, biophysical setting, and stand conditions.

## Methods

### Study Site

Data were collected in the Monument Canyon Research Natural Area (RNA), in the Jemez Mountains of north-central New Mexico, USA (35.8° N, 106.6° W). The site was established as one of the earliest RNAs in the region (1935), and was thus protected from land uses such as logging or livestock grazing (Swetnam et al. 2015). Fire was excluded from the RNA and the surrounding landscape for most of the 20^th^ century, which represents a significant deviation from the historical regime of high frequency-low intensity fire (Falk 2006, Falk et al. 2007). An 8×9 grid of 0.25-ha 50×50 m plots, centered 500 m apart, was established in 1999 and 2000 (described in Falk 2004, Farris et al. 2013). Fifteen of these plots were included in the present study, ranging in elevation from 2520-2590 m, with forest types ranging from essentially pure *Pinus ponderosa* to dry mixed conifer stands. These plots vary with respect to substrate, slope, aspect (Table 1), and other variables. Twentieth century fire suppression led to overly dense forest, particularly on pumice-derived soils, such that a combined thinning (2006) and prescribed fire (2012) treatment was applied to restore forest structure. Our sample included 3 plots that experienced these treatments, one on each of the three substrate types present (tuff, pumice, and alluvium; Table 1). The site is characterized by cool winters and warm summers (mean January and July temperatures for the period 1982-2013 were −2.1°C and 17.9°C, respectively), with wet winters (mean November-March total precipitation = 202 mm) and peak precipitation in July and August (mean total precipitation = 164 mm) associated with the North American monsoon (Figure 1).

**Figure 1.**
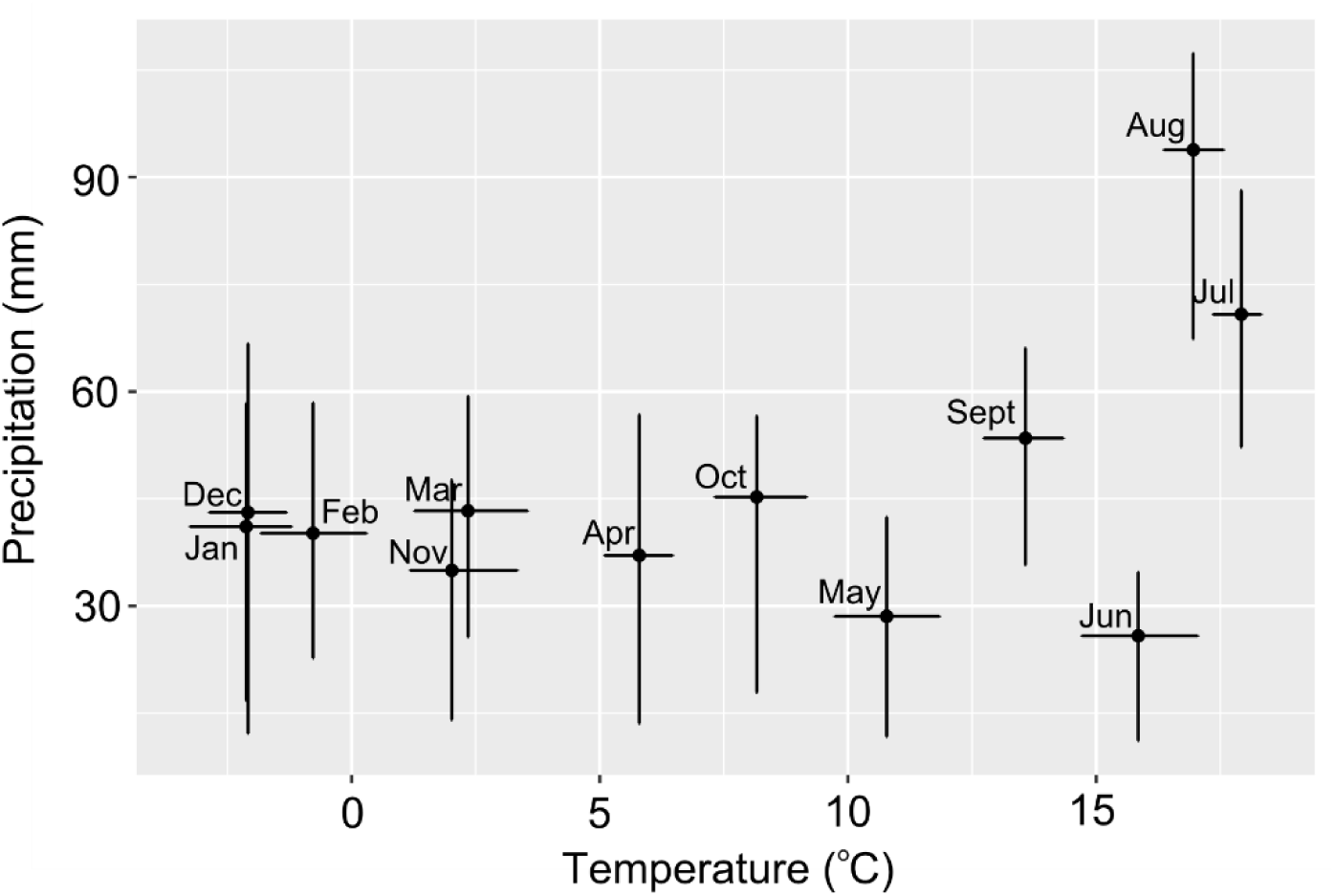
Climate at the Monument Canyon Research Natural Area, Jemez Mountains, New Mexico, U. S. A. (35.8° N, 106.6° W). Data are the 1982-2013 4-km resolution PRISM time series of monthly mean temperature and total precipitation. Bars indicate the 25%-75% quantile of the data.

**Table 1.** Plot-level information at the Monument Canyon Research Natural Area study site, including plot number, elevation, covariates used in the model, and sample sizes of the two data types (diameter at breast height [DBH] and annual tree rings). Elevation is in meters (m) and radiation is expressed in watts m^-2^yr^-1^. Aspect is measured in radians from zero (due north) to 6.28 (360°). The wetness index (wet) is derived from total cachment area (m^2^) and the tangent of the slope (unitless; see Boehner et al. 2002). Substrate (subs) is either tuff (T), pumice (P), or alluvium (A), from Kelley et al. (2003). Basal area (BA, in the year of the first census, either 1999 or 2000) is measured in m^2^ha^-1^. Thinning and fire treatments were applied in 2006 and 2012, respectively. Sample sizes shown are the number of trees measured for DBH and annual ring increments.

| Plot | elev | rad | slope | aspect | wet | subs | BA (m<sup>2</sup>/h) | Thin | Fire | DBH | rings |
| --- | --- | --- | --- | --- | --- | --- | --- | --- | --- | --- | --- |
| 113 | 2541 | 3561 | 0.17 | 3.05 | 6.98 | T | 15.99 | 1 | 1 | 25 | 18 |
| 125 | 2565 | 3407 | 0.16 | 2.55 | 6.80 | P | 28.56 | 1 | 1 | 34 | 12 |
| 127 | 2520 | 2942 | 0.11 | 2.967 | 10.13 | T | 56.97 | 0 | 0 | 80 | 8 |
| 137 | 2564 | 3993 | 0.16 | 5.34 | 7.36 | T | 22.99 | 0 | 0 | 36 | 6 |
| 141 | 2521 | 3666 | 0.07 | 3.15 | 9.80 | A | 27.37 | 1 | 1 | 36 | 10 |
| 143 | 2566 | 3760 | 0.06 | 2.34 | 7.83 | P | 13.43 | 0 | 0 | 7 | 4 |
| 145 | 2533 | 4018 | 0.12 | 2.93 | 7.49 | T | 28.66 | 0 | 0 | 35 | 15 |
| 147 | 2545 | 3404 | 0.18 | 5.69 | 8.55 | A | 17.92 | 0 | 0 | 28 | 11 |
| 149 | 2586 | 3386 | 0.09 | 3.63 | 9.18 | P | 12.89 | 0 | 0 | 13 | 9 |
| 151 | 2532 | 2268 | 0.12 | 2.35 | 7.64 | T | 34.11 | 0 | 0 | 29 | 10 |
| 156 | 2555 | 2891 | 0.34 | 2.41 | 7.28 | T | 29.49 | 0 | 0 | 37 | 10 |
| 158 | 2589 | 3972 | 0.19 | 2.27 | 7.81 | P | 43.13 | 0 | 0 | 34 | 8 |
| 162 | 2537 | 3361 | 0.25 | 2.38 | 7.42 | T | 28.40 | 0 | 0 | 38 | 8 |
| 164 | 2556 | 2979 | 0.15 | 5.79 | 8.96 | T | 28.27 | 0 | 0 | 39 | 12 |
| 166 | 2585 | 3674 | 0.20 | 4.70 | 8.82 | T | 24.69 | 0 | 0 | 38 | 7 |

### Growth increment data

At the time of plot establishment, all individuals of *Pinus ponderosa* >25 cm diameter at breast height (DBH) were tagged and measured (>20 cm DBH threshold for all other species, including *Abies concolor*, *Pinus flexilis*, and *Pseudotsuga menziesii*). In a 10×10 m subplot, all trees (regardless of size) were tagged and measured at breast height. In February and April of 2015, a second measurement of DBH, and first measurement of height (in m, by laser hypsometer) was collected from every tagged tree, thus capturing growth up to the end of the 2014 growth season (and not the 2015 growth season). Short increment cores were collected from ~10 trees per plot, randomly chosen within four size classes (0-20 cm, 20-35 cm, 35-50 cm, and 50+ cm DBH), irrespective of species identity. These samples were prepared following standard dendrochronological methods (Speer 2010). They were then crossdated visually referencing a chronology developed at Banco Bonito (Farella 2015); annual ring dating accuracy was verified using the program COFECHA (Holmes 1983). Annual ring widths were then measured to the nearest μm, for the period 1950 to 2014, on a sliding stage micrometer. The majority of the data were derived from samples of *Pinus ponderosa* (87% of increment cores, 81% of DBH measurements), with limited representation of *Abies concolor*, *Pinus flexilis*, and *Pseudotsuga menziesii* (7%, 5%, 1% of increment cores and 10%, 4%, and 5% of DBH measurements, respectively).

### Covariate Data

Tree-level predictors of radial and diameter growth increments included size (basal area at the time of the first DBH measurement, either 1999 or 2000) and height ratio (the ratio of a given tree’s height compared to the tallest tree on the same plot in 2015). Plot-level predictors included plot basal area at the time of the first DBH measurement, disturbance history (thinning and prescribed fire; Table 1), and several GIS-derived biophysical variables (substrate, slope, aspect, annual radiation, and a wetness index following Boehner et al. 2002). Plot basal area was calculated by converting DBH data for each tree to basal area 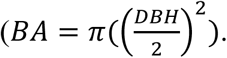 Total BA of large trees (those above the 20 cm DBH threshold for all species but *Pinus ponderosa*, above the 25 cm DBH threshold for *Pinus ponderosa*, censused in the 50×50 plot) was multiplied by 4 to convert to a per-hectare basis. Total BA of small trees (below species-specific size thresholds, censused in the 10×10m inner plot) was multiplied by 100 to convert to a per-hectare basis. The two were summed and expressed in units of m^2^ ha^-1^. Plot basal area ranged from 13.43 to 56.93 m^2^ ha^-1^ among the 15 plots (Table 1). Time series of monthly mean and maximum temperature (°C), and precipitation (mm) were derived from 4 km-resolution AN81m PRISM data (Jan 1982-Dec 2013; PRISM Climate Group, Oregon State University, http://prism.oregonstate.edu). All plots had the same climate data; *i.e*., they all fall in the same PRISM cell. Vapor pressure deficit (VPD) data were derived from Climate Research Unit (CRU) 3.21 products, as saturated minus actual vapor pressure (in hectopascals). All covariate data were centered and scaled (to a mean of zero and a standard deviation of 1.0), so that the magnitude of their effects could be compared.

### Dendroclimatic Analysis

We conducted a dendroclimatic analysis of the tree-ring data, for comparison against the Bayesian multiple regression approach. In tree-ring studies, low-frequency variation in the absolute value of radial increments are removed (*i.e*., the long-term reduction in radial increment that results from increasing diameter over time), a procedure known as “detrending” (Speer 2010). We used a cubic smoothing spline with a 50% frequency cutoff response at 10 years to detrend the tree-level measurement series (function detrend, R package dplR, Bunn 2008, 2010). This retained annual to decadal growth variability in the data while removing longer-term trends. A pre-whitened, site-level mean chronology was calculated from the total pool of 148 detrended series using a bi-weight robust mean (function chron {dlpR}; Bunn 2008). Monthly correlation and response functions were calculated using the function dcc in the R package treeclim (Zang and Biondi 2015). Correlations were pairwise (univariate) Pearson’s correlations between the site-level chronology and monthly PRISM climate data spanning a 16-month window from the previous year’s May to the current year’s August. Response functions use a principal components regression approach (Biondi and Waikul 2004, Zang and Biondi 2015) to examine drivers of tree-ring index variation while accounting for collinearity among climate variables.

### Bayesian fusion

The hierarchical model comprised two regressions, predicting radial increment and diameter increment, respectively, constrained via a constant of proportionality between regression parameters on fixed effects common to both regressions. Motivating this design is the constant of proportionality between the two data types: two times the radial increment is the diameter increment in a single year (setting aside sources of error in both measurements). Thus we expect a constant of proportionality between (for example) the effect of plot-level radiation on annual radial increment vs. 10-year diameter increment. The shared fixed effects linking the two regressions together included two tree-level variables (tree size and height ratio) and five plot-level variables (radiation, slope, aspect, wetness index, and plot basal area; Table 2). In the following, we detail each regression in turn, and then the connection between them.

**Table 2.** Predictors included in the Bayesian multiple regression mixed effects model. Each variable is described as either a fixed or random effect, with respect to the sampling unit to which it applies, and as either continuous or an indicator. Tree size (basal area) and plot basal area, as predictors of growth increment, were measured at the time of the first census. Height ratio was based on heights exclusively measured at the time of the second census. Tree-ring and DBH columns indicate (X) which variables enter in the two submodels, respectively, and the column “coupled” indicates (with an α) those effects whose estimates were coupled to one another across the two submodels via a constant of proportionality.

| Variable | Fixed vs.<br>Random | Indexing | Scale | Tree-ring | DBH | coupled |
| --- | --- | --- | --- | --- | --- | --- |
| Tree size (basal area) | Fixed | Tree | Continuous | X | X | $\alpha$ |
| Height ratio | Fixed | Tree | Continuous | X | X | $\alpha$ |
| Plot basal area | Fixed | Plot | Continuous | X | X | $\alpha$ |
| Radiation | Fixed | Plot | Continuous | X | X | $\alpha$ |
| Slope | Fixed | Plot | Continuous | X | X | $\alpha$ |
| Aspect | Fixed | Plot | Continuous | X | X | $\alpha$ |
| Wetness Index | Fixed | Plot | Continuous | X | X | $\alpha$ |
| Thinning + Fire | Fixed | Plot | Indicator | -- | X | -- |
| PRISM climate data | Fixed | Year | Continuous | X | -- | -- |
| Plot ID | Random | Plot | Indicator | X | X | -- |
| Tree ID | Random | Tree | Indicator | X | -- | -- |
| Species ID | Random | Species | Indicator | -- | X | -- |

The tree-ring submodel was a multivariate normal Gaussian process model, which accounts explicitly for autocorrelation in the residuals *є_t_*, *i.e*., lag effects in the time series of radial increments (*r.inc*) caused by, for example, physiological carry-over in resources that affect tree growth. Two tree-level predictors were included (according to the individual tree index *i*): tree size (*S*, basal area at the time of the first DBH census) and height ratio (*HR*; Table 2). Plot-level fixed effects were radiation, slope, aspect, wetness index, and plot basal area (*rad*, *slope*, *asp*, *wet*, and *PBA*, respectively, using the plot index *j*). Because monthly mean vs. maximum temperature vs. vapor pressure deficit were strongly collinear (Supplementary Figure 1a), and tree responses to these variables were similar (compare Figure 2a to Supplementary Figure 1b), we included monthly mean temperatures (vector ***T***, with the year index *t*), along with monthly total precipitation (vector ***P***), as predictors in the model. Plot random effects (*є*. *p_j_*), normally distributed with a prior mean of zero and variance *σ*.*p^2^*(the latter estimated from the data), were included to account for variation not captured by the plot-level fixed effects, for example, substrate that is known to vary among the plots (Table 1). Individual random effects (*є*.*i_i_*), normally distributed with a prior mean of zero and variance *σ*,*i^2^*, were also included. Residual variation (*є_t_*) was modeled as multivariate normally-distributed, with a mean of zero and variance-covariance matrix Σ*_mn_*, where *m* and *n* refer to years, each ranging from one to 32, thus Σ*_mn_* has dimensions 32×32. The full tree-ring (*r.inc_ijt_*) submodel is expressed as:

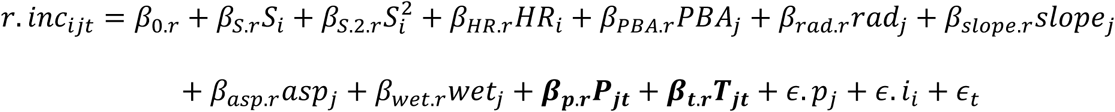

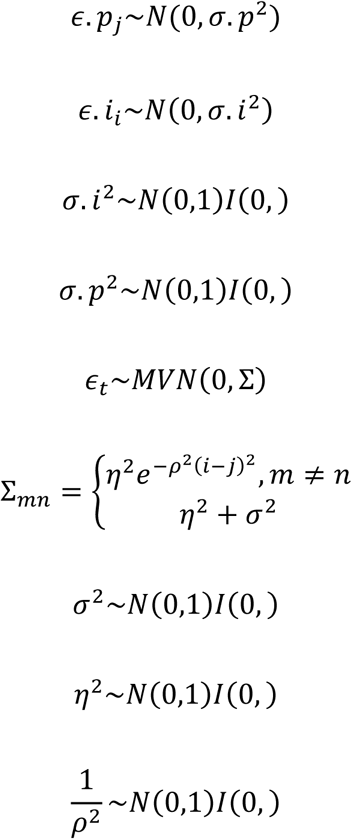

**Figure 2.**
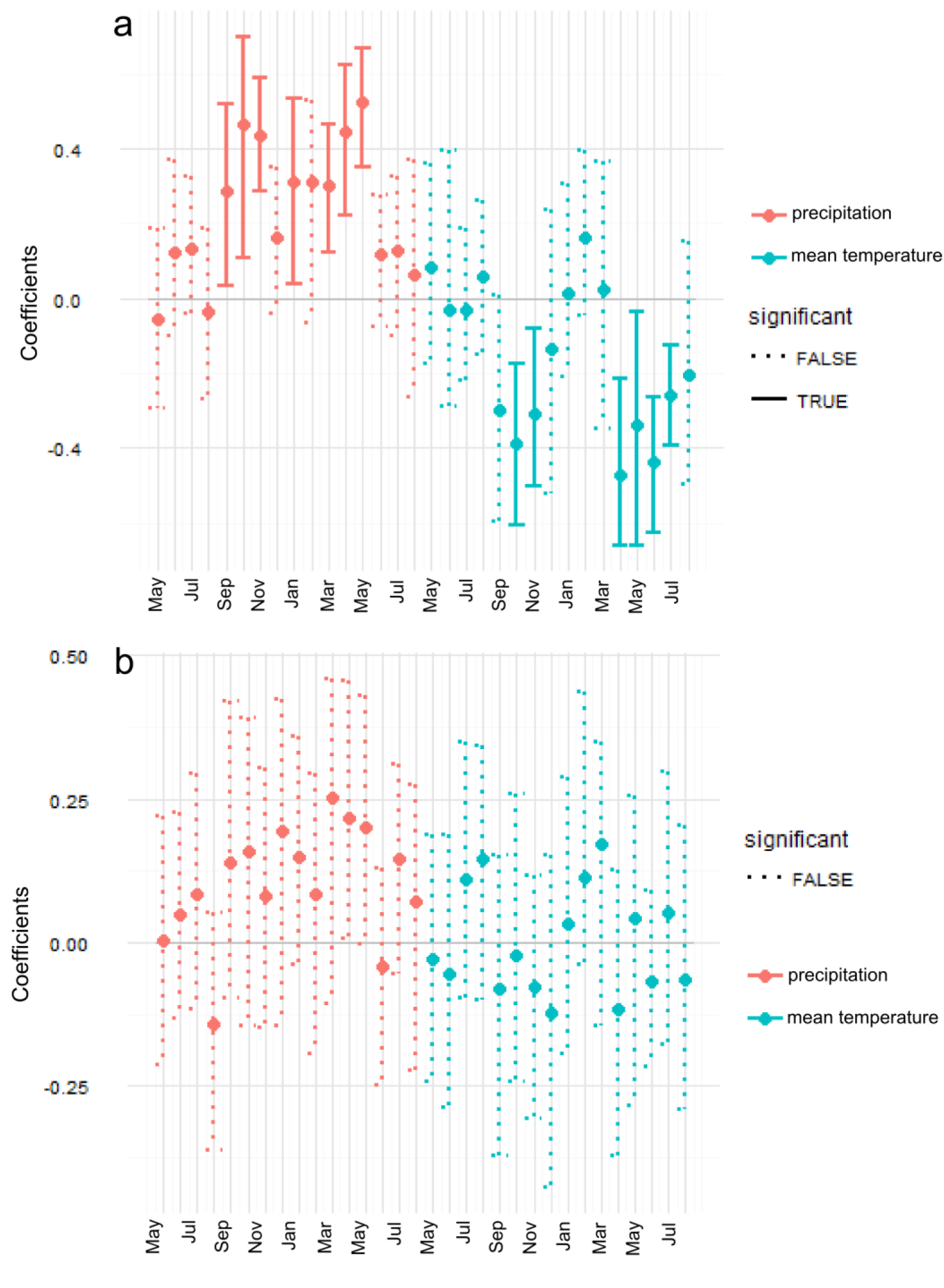
Climate correlation function (a) and response function (b), based on a single, site-level tree-ring chronology and PRISM 4-km resolution monthly total precipitation and mean temperature time series data (1981-2014).

Where the parameter *η* is within-year residual variation, and the parameter *ρ* is the rate at which covariance (among years) decays.

The diameter increment (*d.inc_ijkt_*) submodel assumes a normally-distributed response, with the same tree-level and plot-level fixed and random effects as above, with the exception that thinning+prescribed fire (*thin.fire_j_*) was included as a plot-level indicator variable (Table 2), there were no individual random effects, and there were species random effects (normally distributed with a prior mean of zero and variance *σ.s^2^*, indicated by the index *k*). No climate predictors were included, since 4 km-resolution PRISM climate data did not vary among the plots, and there was only one measurement of diameter increment per tree.

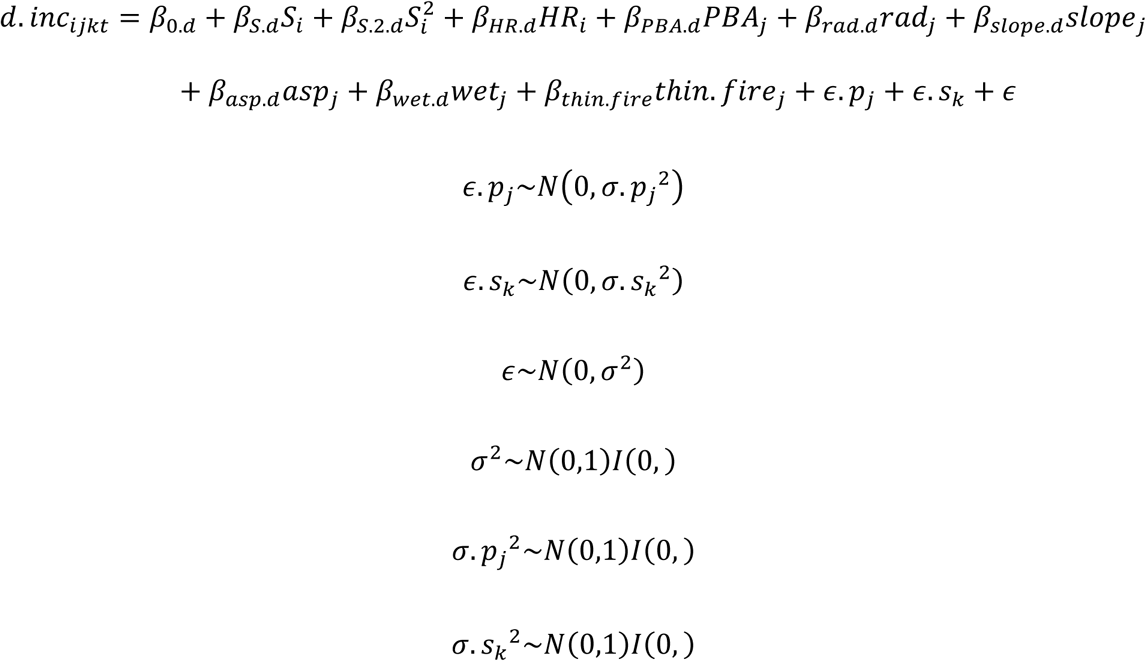

The two regressions were coupled via a constant of proportionality (*α*) between shared fixed effects:

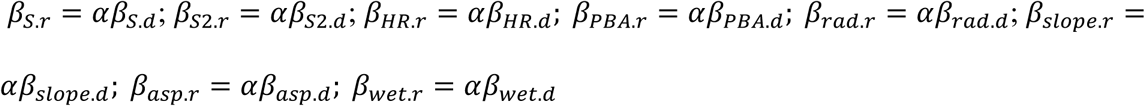

Prior distributions for the regression coefficients were normal with mean of zero and variance of 1.0. The prior for *α* was a positive half-normal, with a prior standard deviation of 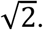 The model was built and executed in R and stan (package rstan; Gelman et al. 2015). Markov chain Monte Carlo (MCMC) simulations consisted of four chains run for 2500 iterations, with the first 1250 steps removed as burn-in. MCMC output was not thinned, leading to 5,000 post-burn-in samples per model parameter. Convergence among chains was evaluated using the *R̂* statistic (Gelman and Rubin 1992); for all model parameters, 0.99 > *R̂* < 1.01, indicating good mixing of the chains.

To compare our multiple regression model (which does not detrend the radial increments) against the dendroclimatic analysis, we first analyzed the radial and diameter increment data using a model that included monthly climate variables (vectors in Equation 1 above). Climate data in a 12-month window from September of the previous year to August of the current year were used to predict tree growth, because the Gaussian process model of radial increments accounts for lag effects caused by climatic conditions in years t-1, t-2, *etc*. Correlations among the 24 monthly climate variables (12 months of mean temperature and precipitation) were generally low (Supplemental Figure 2a); only 6% of pairwise Pearson’s correlations exceed an absolute value of 0.50 (excluding correlations of 1.0 on the main diagonal of the correlation matrix), suggesting that multicollinearity should not affect model stability. Principal components analysis of the monthly climate time series data shows little clustering, with only 42.2% of the total variance explained by the first 3 principle component axes (Supplemental Figure 2b).

We then formed a second model of reduced complexity – i.e., four seasonally aggregated climate variables – and investigated interaction effects of *a priori* interest. The seasonal variables were formed based on the dendroclimatic analysis, which identified climate in certain seasons as limiting tree growth (see Results), and the literature suggesting a general climate response of tree growth across the southwestern and interior western U. S. (Littell et al. 2008, Chen et al. 2010, Williams et al. 2013, Dannenberg and Wise 2016). The four seasonal variables were warm season (previous September and October plus the current April-August) mean temperature and cumulative precipitation and cool season (previous November to current March) mean temperature and cumulative precipitation. The warm season months are when drought stress is most likely to occur, whereas the cool season months are when repeated frontal storms promote deep soil infiltration, fueling early season tree growth (citations). These seasonal variables are significantly correlated (Supplemental Figure 2c), although below the threshold value of |r|= 0.70 that is commonly considered problematic (Dormann et al. 2013). To verify model stability (in the face of collinearity between seasonal climate variables), we formed a variety of simpler vs. more complex multiple regression models, as well as splitting the data in half (randomly) and running the model on each half of the data, and examined the stability of parameter estimates.

In this second model, we tested the interaction between plot basal area and climate, *i.e*., two forms of stress, competition and climate, that are expected to exacerbate one another, such that individual tree growth should be more greatly reduced by climate stress (high temperature, low precipitation) if a tree is in a high density stand than a low density stand. We also tested interactions between climate and height ratio (*i.e*., tree status in the canopy, relative to neighbors), and between climate and tree size (basal area at the time of the first census).

### Model comparison

To evaluate whether a model that combines tree-ring and DBH data together outperforms separate models, we used a leave-one-out (LOO) cross-validation procedure. The principle of LOO cross-validation is to evaluate the log-likelihood of a single data point that was not included in parameter estimation, and repeat this for each data point, yielding an estimate of overall model out-of-sample predictive accuracy. LOO log-likelihoods of the tree-ring and DBH data were compared between the model described above, in which tree-ring and DBH submodels were coupled via the constant of proportionality α (“coupled”), vs. the case where they were uncoupled from one another (*i.e*., separate regressions; “uncoupled”). Similar to Akaike’s information criterion (AIC), a lower expected log pointwise predictive density (elpd) indicates better predictive accuracy; the standard error of the difference in elpd between two models can be used to assess whether one model has demonstrably better out-of-sample predictive performance than another (Vehtari et al. 2016). This LOO comparison was made using the R package loo, which implements Pareto smoothed importance sampling, and conveniently takes as input MCMC simulation draws from a stan model (Vehtari et al. 2016).

### Reduced Data Scenario

The dataset we developed was exceptionally rich with respect to increment cores (129 in a small area). In order to better gauge the value of combining the two data sources (tree-ring and DBH remeasurement), we created a more realistic data scenario in which the number of increment cores was dwarfed by forest inventory-type data. We randomly sampled a single increment core from each of the 15 plots, then reran the coupled and uncoupled models (“model 1”, with 24 climate variables and no interaction effects), and evaluated a) the performance of the coupled vs. uncoupled models (in terms of LOOIC) and b) the posterior distributions of effects.

## Results

### Dendroclimatic Analysis

Detrended radial increments were negatively correlated with temperature in the preceding September-November and current year April-July, and positively correlated with precipitation in similar months (Figure 2a), indicating that tree growth is precipitation-limited at this site. This is a typical climate signal for tree growth in the southwestern U. S. (Williams et al. 2013). Correlations between growth and a) maximum temperature or b) vapor pressure deficit were very similar to the correlations with mean temperature (compare Supplementary Figure 1b to Figure 2a). The response function also indicates that growth is positively sensitive to precipitation and (to a lesser degree) negatively sensitive to temperature, although these effects are not significant based on bootstrapping of a 30-year time series (Figure 2b).

### Bayesian fusion

The Bayesian multiple regression model that included 24 monthly climate variables inferred univariate associations very similar to the dendroclimatic analysis: positive effects of precipitation and negative effects of temperature, especially at the end of the previous year’s growing season and the beginning of the current year’s growing season (compare Figure 3a to Figure 2a). Partial regression coefficients, which account for the effects of the other 23 climate variables and correlations with them (Figure 3b) were similar to the dendroclimatic response function (Figure 2b) in that they consistently showed strong positive effects of precipitation vs. weaker effects of temperature, including both negative and positive temperature effects.

**Figure 3.**
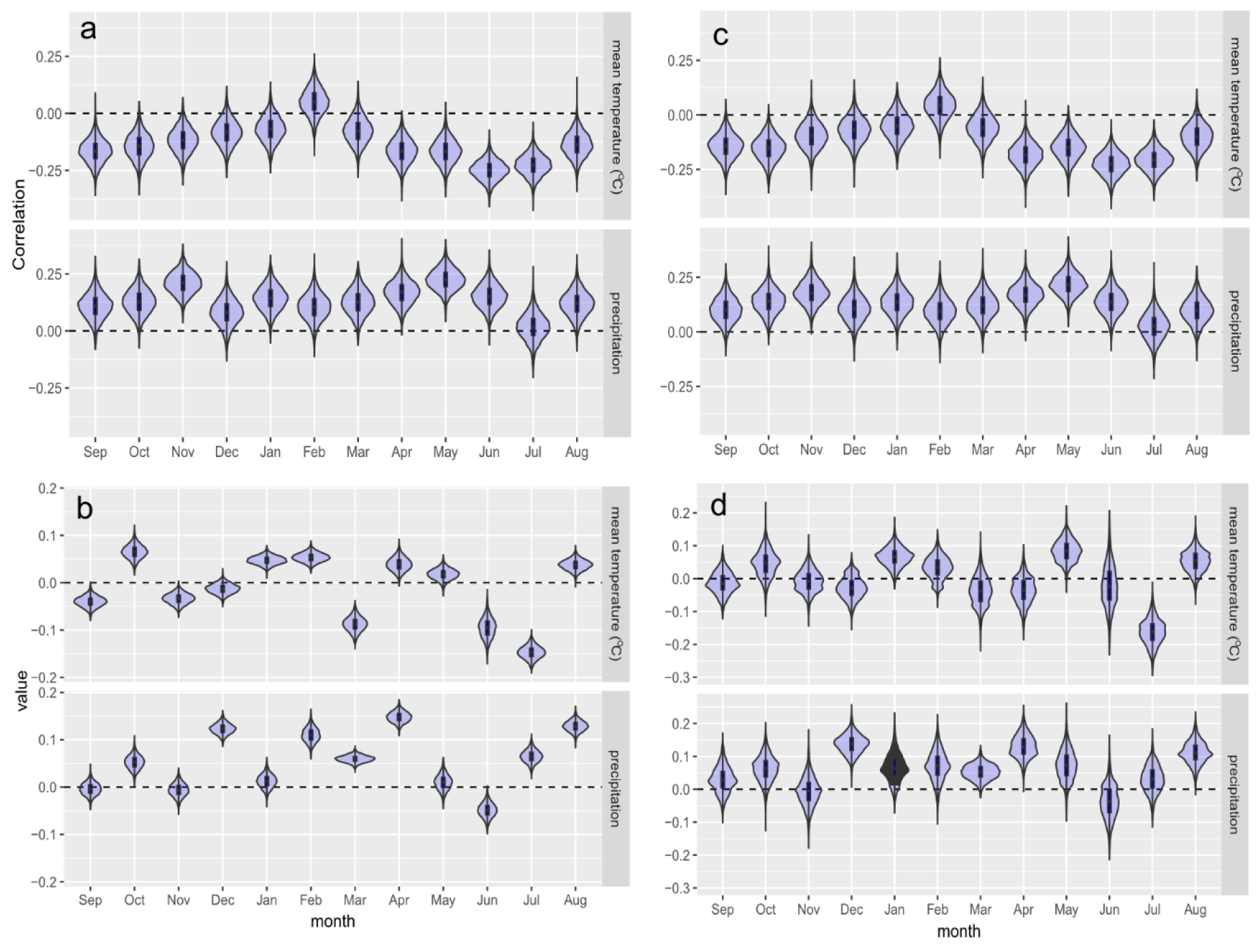
Posterior distributions of climate effects from the “uncoupled” version of the Bayesian multiple regression model. (a) Bivariate associations, calculated from partial regression coefficients and correlations between each variable and the other 23 monthly climate variables and their effects. (b) Partial regression coefficients. (c) Bivariate associations and (d) partial regression coefficients under the reduced data scenario of a single increment core randomly selected from each plot (n=15). Note that the y-scale in panel d is larger than in panel b.

Of all the factors in this model, including the 24 climate variables, tree size had the strongest effect on radial and diameter increments: growth increments declined with tree size, but this slowed with increasing size (negative and positive first-order and second-order terms, respectively; Figure 4). Height ratio had a positive effect on growth increments, and plot basal area had a negative effect (Figure 4). Aspect had a negative effect on growth, as did radiation, whereas the wetness index had a positive effect (Figure 4). The effect of slope on diameter and radial increments was indistinguishable from zero. The effect of canopy removal (thinning followed by fire) on diameter increments was relatively large (the second largest effect by magnitude, after the effect of size), but the posterior distribution of this effect overlaps zero considerably. Investigation of the interaction between thinning+fire and substrate, via plot random effects, showed that growth was somewhat elevated in plots on a) pumice and alluvium soils that experienced thinning and prescribed fire, compared to b) thinned and burned plots on tuff soils and c) plots that were not thinned and burned (Figure 5), consistent with the fact that Pueblo farmers in the area are known to have focused their attention on pumice soils for their productivity (Gauthier et al 2007).

**Figure 4.**
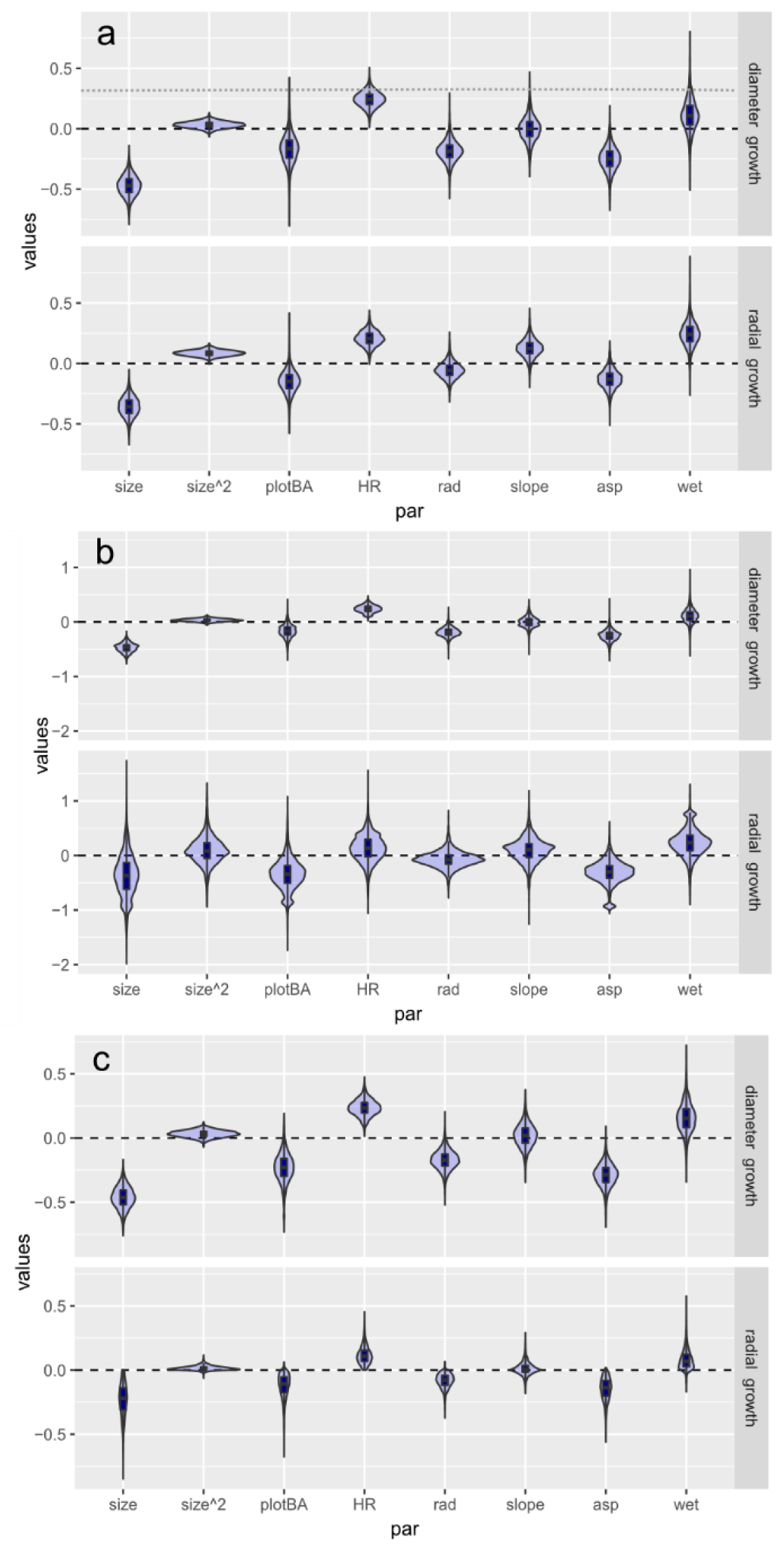
Posterior distributions of non-climatic fixed effects on diameter growth (DBH submodel) vs. radial growth (tree-ring submodel), including individual tree size and the quadratic size term (size^2^), plot basal area, height ratio (HR), radiation, slope, aspect, and wetness index.(a) Uncoupled model, using 500 remeasurements of DBH and 129 increment cores. A second horizontal line in the top panel indicates the posterior mean effect of thinning followed by fire.(b) Uncoupled version, using just 15 increment cores (note increased y-scale). (c) Coupled model, using 15 increment cores.

**Figure 5.**
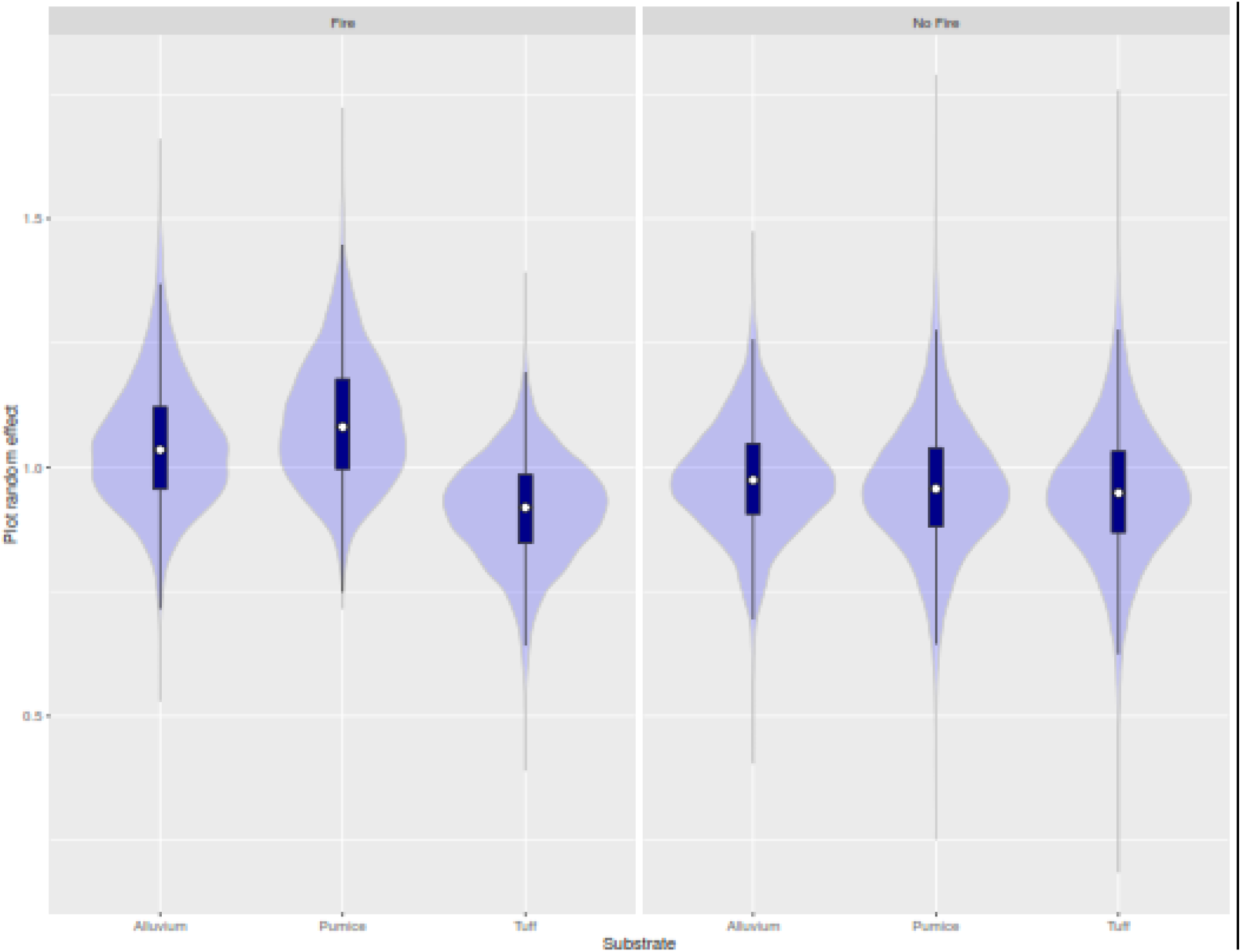
Posterior distributions of plot random effects, grouped according to whether the plot was thinned (2006) then burned (2012) vs. substrate (alluvium, pumice, or tuff).

Seasonally-defined precipitation effects on growth were positive and temperature effects negative. Cool and warm season precipitation had the strongest effects on tree growth. Warm season temperature had a weaker but non-zero effect on growth. Cool season temperature did not affect growth. Other main effects were as before: size and size^2^ had negative and positive effects, respectively, height ratio had a positive effect, plot basal area had a negative effect, aspect and radiation had negative effects, wetness had a positive effect, and slope did not consistently affect growth (the effect of thinning+fire was not tested, since this effect proved indistinguishable from zero in the model with 24 monthly climate variables). Interaction effects between seasonal climate variables and tree size, plot basal area, as well as height ratio were indistinguishable from zero.

### Model Comparison

The difference in LOO log-likelihoods of radial increments between the coupled and uncoupled models was 0.31 (standard error of 2.41), indicating that the out-of-sample predictive performance of the two models was indistinguishable with these data (model 1, with 24 climate variables and no interaction effects). The same was true with respect to the diameter increments: the difference between LOO log-likelihoods (coupled-uncoupled) was 0.11 (se = 1.62).

### Reduced Data Scenario

The coupled and uncoupled models were also statistically indistinguishable from one another when only 15 increment cores were analyzed (rather than 129): the difference between LOO log-likelihoods (coupled-uncoupled) was 0.38 (se = 0.65) for the radial increments (tree-ring data) and −0.15 (se = 0.52) for the diameter increments (DBH data). Posterior distributions of climate effects (partial regression coefficients) were noticeably broader under the reduced data scenario (Figure 3d), as expected, since only the tree-ring data inform estimates of climate effects. The same was true of non-climate fixed effects estimated from the tree-ring data, under the uncoupled model (Figure 4b). In contrast, under the coupled model, posterior distributions of non-climate effects on radial growth were similar (mean and variance) to what was obtained using the much larger sample of increment cores (Figure 4c).

## Discussion

The Bayesian hierarchical model that we present is capable of taking advantage of the strengths of two widely available and complementary data sources – tree-ring and forest inventory - to estimate the influence of many factors on tree growth. That is, the model formally combines the two data sources, and accommodates the complexity of individual tree growth, including influences at multiple scales (tree-level and stand-level) that jointly result in variation in growth among individual trees. The results from this proof-of-concept analysis confirm several expectations about tree growth at a single, well-studied site. That is, the primary influence on growth increments is tree size, followed by a negative effect of plot basal area and positive effect of height ratio, as well as a suite of effects consistent with precipitation-limited tree growth – negative effects of radiation, aspect, and average temperature, combined with positive effects of a wetness index and cumulative precipitation (Figures 3 and 4).

That tree size had the strongest effect on radial and diameter growth increments is not surprising – this is, first and foremost, the effect of geometry. Given a certain amount of photosynthate (biomass) produced by a tree, with increasing basal area each year, the new biomass is spread out over a larger and larger circumference. The result is a negative effect of basal area on radial and diameter increments, weakening with increasing size (negative and positive first-order and second-order terms, respectively; Figure 4). Tree-ring data have traditionally been detrended (or “standardized”) to remove low-frequency variation in radial increments caused by changing tree size or, importantly, changes in stand conditions (an issue we return to below). This practice is problematic because it removes low-frequency variation regardless of the cause, be it the increasing size of the tree or long-term trends in temperature (or other global change factors). We offer an alternative approach to modeling tree-ring data, noting that it depends on knowledge of the absolute size of the tree (*e.g*., DBH) for at least one point in time (e.g., when the increment core was collected), and preferably, also stand basal area or some other measure of stocking density.

Climatic influences on growth detected by the Bayesian multiple regression model were very similar to results from a dendroclimatic analysis, in terms of tree response and its seasonality (Figure 2a vs. Figure 3a). We note that the error bars in Figure 2a are not directly comparable to error bars in Figure 3 a, because the former was created by bootstrapping a 30-year time series of climate data with respect to a single site-level chronology (Zang and Biondi 2015), whereas the latter indicate Bayesian credible intervals of fixed effects in a multiple regression model (which includes factors not in the dendroclimatic analysis: non-climatic fixed effects, plot random effects, *etc*.). The results from these two approaches were also broadly similar with respect to analyses that take into account correlations between climate variables (Figure 2b and 3b), although they differ in details, which is not surprising given that they are derived from different methods (principal components regression of detrended data vs. a hierarchical multiple regression of raw data). When seasonal climate variables were included in the Bayesian model, the magnitude of climate effects were (in decreasing order): precipitation (cool and warm season), followed by warm season temperature. The effect of cool season temperature did not differ from zero.

The estimated effect of thinning followed by prescribed fire on diameter increments was positive and large - a posterior mean of 0.291 (see horizontal line, Figure 4a). But this effect was not distinguishable from zero (standard deviation of 0.217), either because of parameter uncertainty or process variability (or both). Our ability to detect any effect of canopy removal on diameter increments was compromised by the fact that the treatments (thinning in 2006, prescribed fire in 2012) occurred part-way through the remeasurement interval (starting in either year 1999 or 2000 and ending in year 2014); thus only a fraction of the observed diameter increment in a given tree reflects this treatment. Process variability might arise with respect to the effect of prescribed fire, since individual trees can have either a positive or negative response to fire: positive if they are not damaged by fire and fire removes competitors, but negative (at least in the short term) if fire does incur significant damage.

Interaction effects between individual tree size and seasonal climate variables were not significant. Keeping in mind that our sampling design targeted trees above a fairly large size threshold (>25 cm DBH for *Pinus ponderosa*), we still might have expected the largest trees in our sample to be more negatively impacted by climate stress than smaller (but still moderately-sized) trees (McDowell and Allen 2015, Rollinson et al. 2016), but we did not detect such an effect. The empirical literature on interactions between size and climate effects is mixed, with some studies showing significant interaction effects and others not (Chhin et al. 2008, Carnwath et al. 2012, Nehrbass-Ahles et al. 2014). We also would have expected competition and climate stress to interact. Existing literature suggests that thinning can mitigate drought stress (Laurent et al. 2003, Klos et al. 2009, D’Amato et al. 2013, Magruder et al. 2013, Rollinson et al. 2016). We see a need for further investigation of climate sensitivity across tree sizes and stand conditions.

Estimates of effects from the tree-ring data vs. DBH data were consistent with one another, even based on the uncoupled model, in which the parallel fixed effects (β_*S.r*_ VS. β_*S.d*_, β_*HR.r*_ VS. β_*HR.d*_) were free to differ (Figure 4a). That is, the two data sources do not tell different stories about (the sign or magnitude of) effects on tree growth. Tree rings are arguably the better data source, because of their annual resolution and temporal depth, but they are expensive to develop, especially in places where false rings and missing rings make cross-dating an essential part of sample processing. This makes the idea of combing the two data sources appealing, *i.e*., combining few tree-ring data with more forest inventory data. Model comparison showed no advantage to fusing the two data types together, in terms of the leave-one-out information criterion (LOOIC), either when a very large sample of increment cores was used (*n* = 129), or when a much reduced sample size was used (*n* = 15 increment cores). However, there was a clear advantage to the combined use of the two data sets (i.e., the “coupled” model) in terms of posterior estimates of non-climate effects (size, plot basal area, height ratio, radiation, topographic wetness index), when there were few increment cores and many DBH remeasurements. Estimates of the non-climate fixed effects were poor (overlapped zero) from the tree-ring data when the uncoupled model was used on a reduced tree-ring data set (Figure 4b), but when the estimates of effects from the two data sources were coupled to one another, the posterior distributions of non-climate effects were forced to resemble one another (tree-ring vs. DBH data, Figure 4c), thus the coupled model was able to "borrow strength" from the rich DBH data, and reduce uncertainty about shared fixed effects substantially (Figure 4c compared to 4b). The performance of our Bayesian fusion model under a data scenario where strength can be borrowed across tree-ring vs. forest inventory data with respect to climate effects is a target of further investigation.

In this analysis, we treated tree size and proxies for competition (plot basal area, height ratio) as invariant, when in fact we know that they evolve over time. In spite of this, we found reasonable, significant (non-zero) effects of these predictors, likely because we analyzed relatively short time series of growth increment data (32 years). To better take advantage of the long time series provided by tree-ring data, it will be necessary to treat tree size and stand conditions as dynamic. The former problem (changing tree size) is relatively easy to solve, especially if a pith date is known (and see the hidden process model approach of Clark et al. 2007). The second problem, sometimes referred to as the “ghost of competition past” or the “fading record”, is more challenging. It would require the marriage of a model of stand dynamics (e.g., twigs, FVS) with a model of individual tree growth, to properly account for both tree-level and stand-level influences on tree growth across decades of forest development.

As new uses for tree-ring data emerge, beyond the traditional dendroclimatic emphasis, including carbon accounting (Babst et al. 2014a,b,Klesse et al. 2016, Dye et al. 2016), projection of future tree growth (Chen et al. 2010, Williams et al. 2013, Charney et al. 2016), and projection of forest dynamics under climate scenarios (Crookston et al. 2010), the importance of collecting metadata on tree-level and stand-level characteristics with increment cores becomes increasingly clear (Brewer et al. 2010). At present, data in the public repository of tree-ring data (the International Tree-Ring Databank, URL) do not contain such metadata. New tree-ring data networks, such as the U. S. Forest Service’s Interior West-Forest Inventory and Analysis network (DeRose et al. 2016), will enable a new set of questions to be addressed with tree-ring data, especially in conjunction with forest inventory data in the plots where the increment cores were collected.

## Acknowledgements

Our thanks go to Tom and Suzanne Swetnam, David Frank, and Susan Harrelson for facilitating this research. Laura Marshall, Chris Guiterman, Ben Olimpio, Craig Allen, Park Williams, Justin DeRose, and Jeff Oliver provided data or helpful feedback. MEKE acknowledges the support of the College of Science, University of Arizona and USDA-AFRI grant 2016-67003-24944. FB acknowledges funding from the EU Horizon 2020 project "BACI" (grant 640176) and the Swiss National Science Foundation (grant P300P2_154543).

---

**Supplemental Figure 1.**
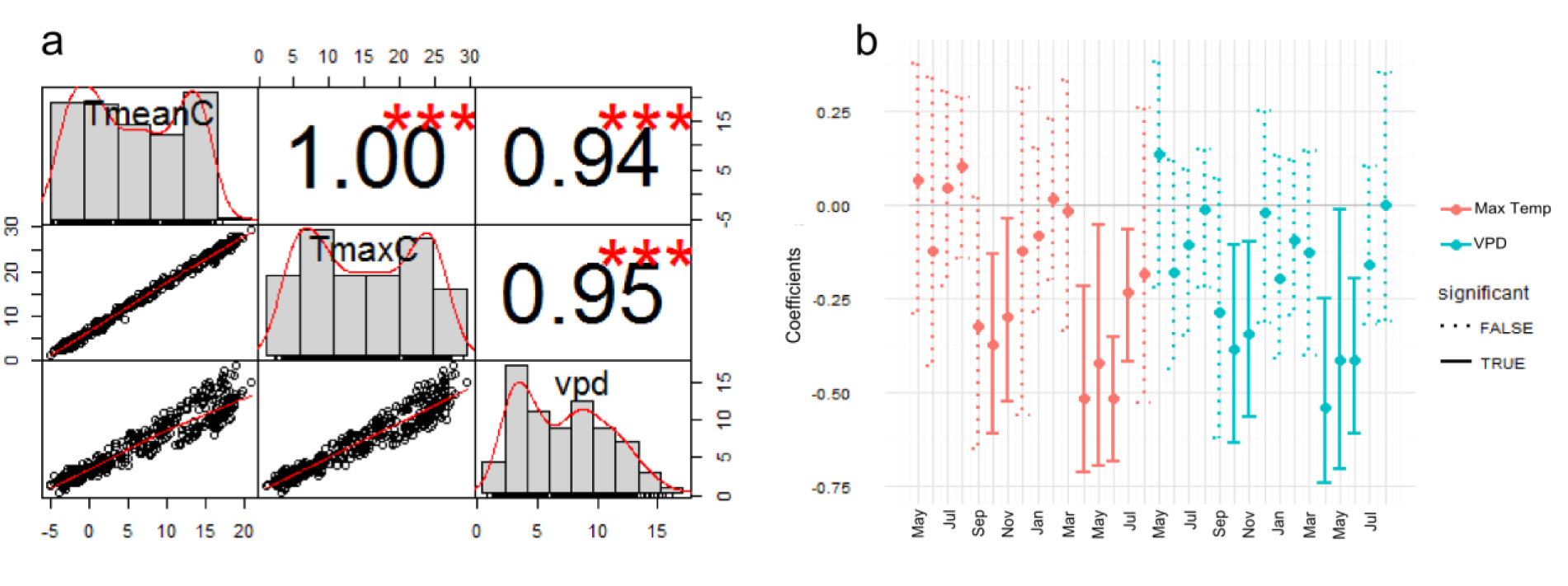
(a) Correlations between three climate variables: PRISM 4-km resolution monthly mean temperature, maximum temperature, and CRU-derived vapor pressure deficit (January 1981-December 2013). Above the diagonal are pairwise correlation coefficients, with asterisks indicating significance. Below the diagonal are bivariate scatterplots of the data, and on the diagonal are frequency histograms showing the distribution of each climate variable.(b) Pearson’s pairwise correlations between (detrended) ring width index and maximum temperature (red) or vapor pressure deficit (blue) are very similar to the correlations with mean temperature in Figure 2a (created via the same use of function dcc{treeclim}; Zang and Biondi 2015).

**Supplemental Figure 2.**
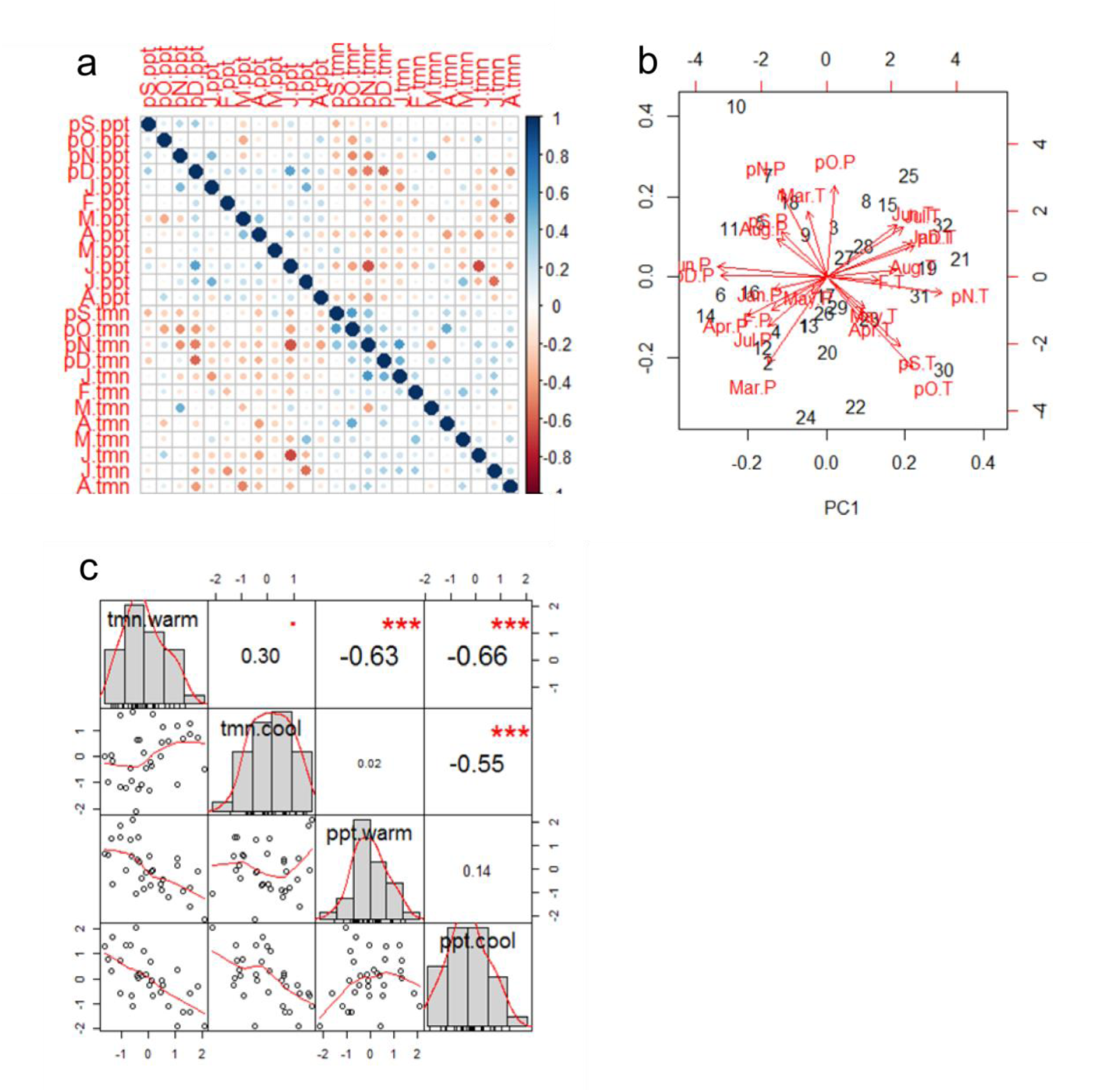
(a) Correlations between 24 monthly climate variables, based on the 4km resolution time series of PRISM monthly total precipitation and mean temperature data, 1982-2013. Climate variables are ordered from previous September (pS) to current year’s August (A), with the additional label indicating precipitation (ppt) or mean temperature (tmn). (b) Clustering of 24 monthly climate variables in principal components space (PC1 vs. PC2, which account for just 32% of the total variance). Numbers are the years from 1982-2013. (c) Correlations between four seasonal climate variables. Above the diagonal are pairwise correlation coefficients, with asterisks indicating significance. Below the diagonal are bivariate scatterplots of the data, and on the diagonal are frequency histograms showing the distribution of each climate variable.

